# Forget Pixels: Adaptive Particle Representation of Fluorescence Microscopy Images

**DOI:** 10.1101/263061

**Authors:** Bevan L. Cheeseman, Ulrik Günther, Mateusz Susik, Krzysztof Gonciarz, Ivo F. Sbalzarini

## Abstract

Modern microscopy modalities create a data deluge with gigabytes of data generated each second, and terabytes per day. Storing and processing these data is a severe bottleneck, not fully alleviated by data compression. We argue that this is because images are processed as regular grids of pixels. To address the root of the problem, we here propose a content-adaptive representation of fluorescence microscopy images called the Adaptive Particle Representation (APR). The APR replaces the regular grid of pixels with particles positioned according to image content. This overcomes storage bottlenecks, as data compression does, but additionally overcomes memory and processing bottlenecks, since the APR can directly be used in processing without going back to pixels. We present the ideas, concepts, and algorithms of the APR and validate them using noisy 3D image data. We show that the APR represents the content of an image while maintaining image quality. We then show that the adaptivity of the APR provides orders of magnitude benefits across a range of image processing tasks. Therefore, the APR provides a simple, extendable, and efficient content-aware representation of images that relaxes current data and processing bottlenecks.

---

Developments in fluorescence microscopy (*1–3*), labeling chemistry (*4*), and genetics (*5*) provide the potential to capture and track biological structures at high resolution in both space and time. Such data are vital for understanding spatiotemporal processes in biology (*6*). Unfortunately, fluorescence microscopes do not directly output the shapes and locations of objects through time. Instead, they produce raw data, potentially terabytes of 3D images (*7*), from which the desired spatiotemporal information must be extracted by image processing. Handling the large image data and extracting information from the raw microscopy images currently presents the main bottleneck (*7–9*). We propose that at the core of the problem is not the amount of *information* contained in the images, but how the *data* encodes this information – usually as a uniform grid of pixels. While data compression can alleviate storage issues, it does not reduce memory usage nor computational cost as all processing must still be done on the original, uncompressed data.

Processing bottlenecks are effectively avoided by the human visual system, which solves a similar problem of inferring object shapes and locations from photon counts. In part, the human visual system achieves this by adaptively sampling the scene depending on its content (*10*), while adjusting to the dynamic range of intensity variations (*11*). This adaptive sampling works by selectively focusing the attention of the eyes on areas with potentially high information content (*10*). This selective focus then enables the efficient inference of information about the scene at a high effective resolution by directing the processing capacity of the retina and the visual cortex. As in fluorescence microscopy, the information in different areas of a scene is not encoded in absolute intensity differences, but in relative differences compared to the local brightness. The human visual system maintains effective adaptive sampling across up to nine orders of magnitude of brightness levels (*11*) by using local gain control mechanisms that adjust to, and account for, changes in the dynamic range of intensity variations. Together, adaptation and local gain control enable the visual system to provide a high rate of information content using as little as 1 MB/s of data from the retina (*12*). In contrast, the information-to-data ratio in pixel representations of fluorescence microscopy images is much lower and is governed by the spatial and temporal resolution of the images rather than by their contents.

Inspired by the adaptive sampling and local gain control of the human visual system, we here propose a novel representation of fluorescence microscopy images: the Adaptive Particle Representation (APR). The APR adaptively resamples an image, guided by local information content, while using effective local gain control. Figure 1A illustrates the basic idea of adaptive sampling. The top panel shows a pixel representation of a fluorescence image acquired from a specimen of *Danio rerio* with labeled cell nuclei. The pixel representation places the same computational and storage costs in areas containing labeled cell nuclei and in areas with only background signal. This uniform sampling results in processing costs that are proportional to the spatial and temporal resolution of the acquisition, rather than the actual information content of the image. A difficulty in adaptation is to give equal importance to imaged structures across a wide range of intensity scales. This is achieved by local gain control as illustrated in Fig. 1B. Without local gain control, adapting effectively to both bright and dim regions in the same image is not possible (Fig. 1B, *centre left*). The APR provides local gain control by guiding the adaptation with a Local Intensity Scale (Fig. 1B, *center right*). As seen in Fig. 1B (*right*), this samples dim and bright objects at comparable resolution, giving them equal importance. Combining adaptive sampling and local gain control, the APR shares two key features of the human visual system to alleviate current processing and storage bottlenecks in fluorescence microscopy.

**Figure 1.**
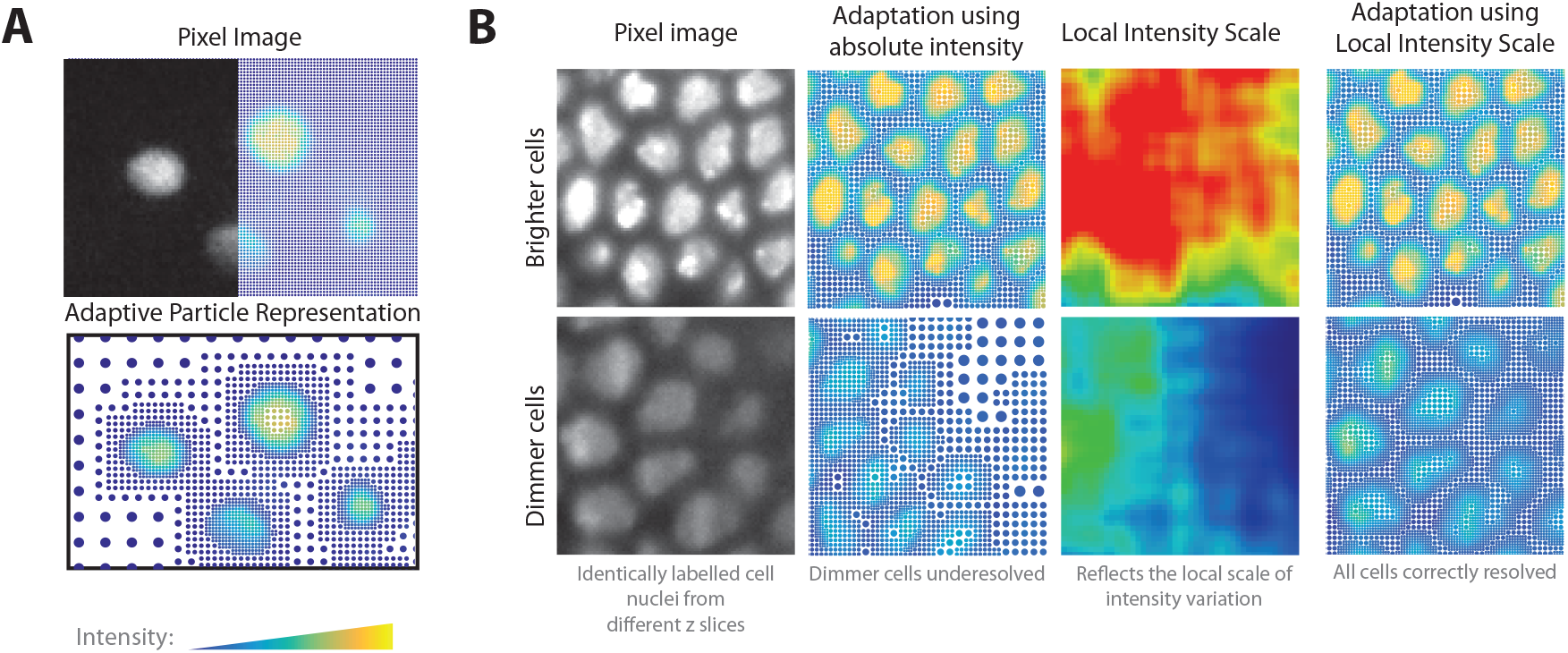
Spatially adaptive representation of fluorescence microscopy images. **A.** Example image of fluorescently labeled zebrafish cell nuclei (Dataset 7 from STable 3, courtesy of Huisken Lab, MPI-CBG & Morgridge Institute for Research), represented on a regular grid of pixels (*top, right half)*. The *top left half* shows the pixel image, and the *bottom* panel the APR. Particles are shown as dots with their color indicating fluorescence intensity and their size reflecting the local resolution of the representation. **B.** Adaptively representing objects of different intensity requires accounting for the local brightness levels. The *left* panel compares two regions of labeled cell nuclei (Dataset 6 from STable 3, courtesy of Tomancak Lab, MPI-CBG) with different brightnesses. The *center left* panel shows adaptive representations based on the absolute intensity. The *right* panel shows the APR accounting for the Local Intensity Scale of the image as shown in the *center right* panel. Using the Local Intensity Scale, objects are correctly resolved across all brightness levels, without over-resolving the background.

While the APR reduces storage costs, as data compression also does, it additionally overcomes memory and processing bottlenecks, since the APR can directly be used in processing without going back to pixels. Compression only alleviates storage costs, as the data need to be uncompressed again for processing or visualization. The APR is therefore not a compression scheme, but an efficient image representation that can additionally then also be compressed. We posit that any image representation aiming to achieve this should fulfil the following *representation criteria* (RC):

RC1: It must guarantee a user-controllable representation accuracy for noise-free images and must not reduce the signal-to-noise ratio of noisy images.
RC2: Memory and computational cost of the representation must be proportional to the information content of an image, and not to its number of pixels.
RC3: It must be possible to rapidly convert a given pixel image into that representation with a computational cost at most proportional to the number of input pixels.
RC4: The representation must reduce both computational cost and memory cost of image-processing tasks with a minimum of algorithmic changes and without resorting to the full original pixel representation.

There is a rich history in multi-resolution and adaptive sampling approaches to image processing, including super-pixels (*13, 14*), wavelet decompositions (*15–17*), scale-space and pyramid representations (*18, 19*), contrast-invariant level-set representations (*20*), dictionary-based sparse representations (*21*), adaptive mesh representations (*22–24*), and dimensionality reduction (*25, 26*). However, none of the existing approaches meets all of the above representation criteria, mainly because they were developed for different applications.

Many previous methods, such as super pixels and contrast-invariant level-set representations, provide effective solutions accounting for changes in spatial scales and contrast. They can efficiently be used for specific tasks, such as image segmentation, providing high-quality solutions at reduced memory and computational costs. However, it is unclear how these methods can be used across a wider range of processing tasks, such as image visualization, without still requiring the original pixel image. Alternatively, adaptive sampling methods, such as thresholded wavelets and adaptive mesh methods, provide more general representations that could replace pixel images while reducing both computational cost and memory cost. However, both approaches have not been adapted to account for local contrast variations are unlikely to be formed rapidly for large 3D images without further improvements. Additionally, techniques that require a change of basis, such as dictionary techniques and wavelets, require the reformulation of image-processing tasks in the transformed domain.

Here, we propose the APR, which meets all of the above representation criteria, and provides a general framework combining concepts from the range of existing methods, resulting in an ideal candidate to replace pixel images in fluorescence microscopy.

## The Adaptive Particle Representation

We first describe the basic concepts of the APR and its components. In the subsequent subsection, we provide all technical details needed to reproduce or reimplement the APR. For simplicity, we do so using a 1D image as a didactic example (see also SuppMat 11; code available from github.com/cheesema/APR_1D_demo). All of the concepts introduced extend to higher dimensions, as shown in the Supplement. Readers not interested in the technical details can skip that subsection and continue to the results and evaluations.

The APR takes an input pixel image and resamples it in a spatially adaptive way that depends on image content, representing it as a set of particles with associated intensity values. Particles are a generalization of pixels, i.e., points in space that carry intensity, but they are not restricted to sit on a regular lattice. Instead, particles can be placed wherever image contents requires, and they may additionally have different sizes in different parts of the image (See Figure 2A).

**Figure 2.**
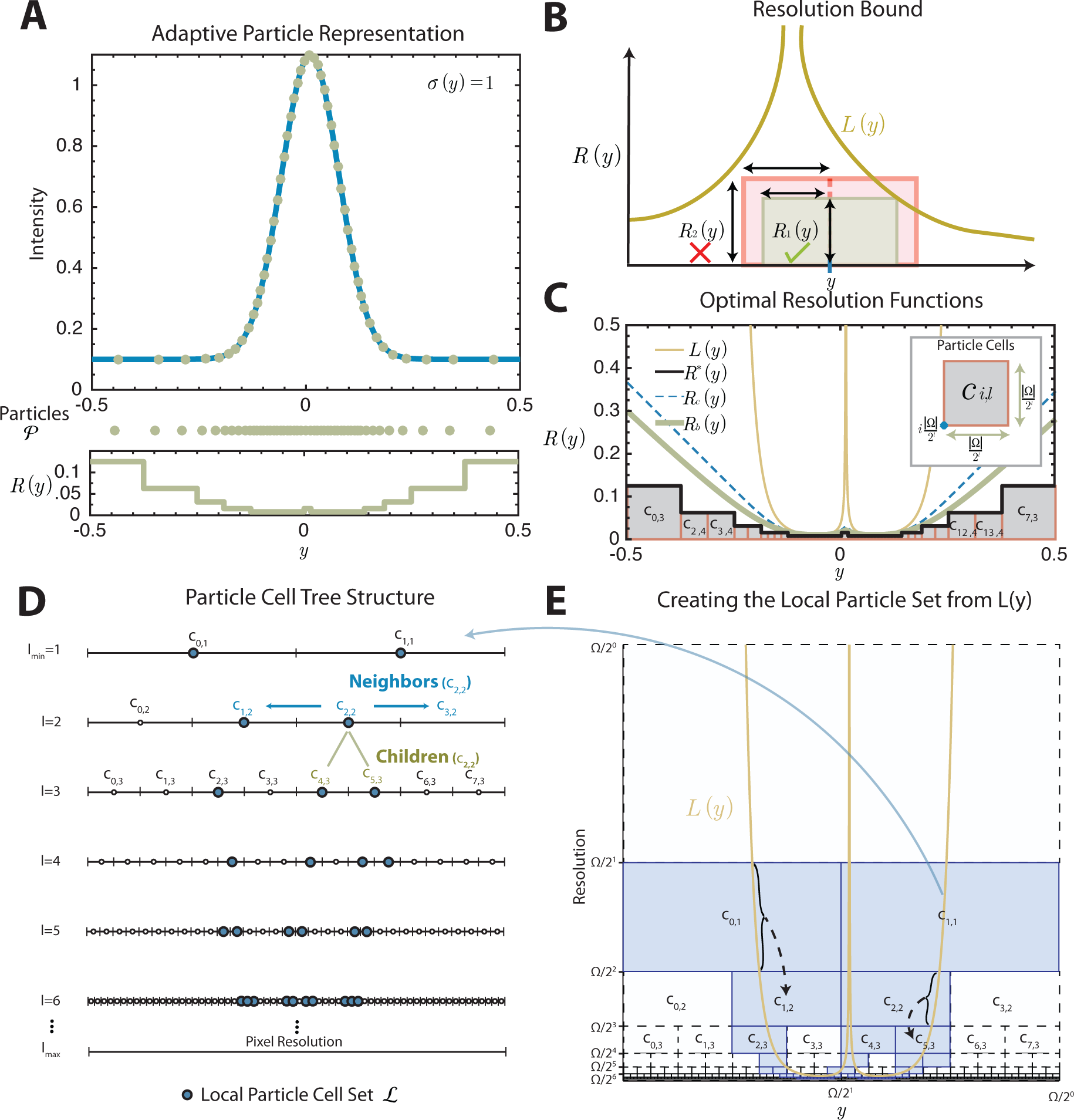
Concepts and definitions of the Adaptive Particle Representation (APR) illustrated in 1D. See main text for explanations. **A.** APR (*E* = 0.1, *σ*(*y*) = 1) representation of the shifted 1D Gaussian 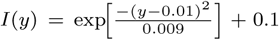. The *bottom* plot shows the corresponding Resolution Function *R*(*y*) with the set of particles shown as dots above. **B.** Illustration of the Resolution Bound, requiring for all pixel locations *y* that a rectangle centered at *y* with width 2*R*(*y*) and height *R*(*y*) does not intersect the graph of the Local Resolution Estimate *L*(*y*). Fulfilling the Resolution Bound guarantees fulfilling the Reconstruction Condition. **C.** Comparison of the optimal (i.e., largest everywhere still satisfying the Reconstruction Condition) Resolution Function *R_c_*(*y*) (blue dashed) with the *R_b_* (*y*) satisfying additionally also the Resolution Bound (bold green) and with the Implied Resolution Function *R^∗^*(*y*) (bold black) for the 1D Gaussian example from (A). The Implied Resolution Function is composed of the upper edges of blocks called Particle Cells (gray). They never intersect the optimal Resolution Function (*R_c_*), therefore providing a conservative approximation. The *inset* shows the Particle Cells definition, described by their level *l* and location *i*. **D.** The set of all possible Particle Cells can be represented as a binary tree reaching down to single-pixel resolution. **E.** The Local Particle Cell set *ℒ* is constructed from *L*(*y*). The correspondences between segments of *L*(*y*) and the Particle Cells in *ℒ* are shown with braces and dashed lines. All possible Particle Cells are shown as blocks, and those belonging to *ℒ* are shaded blue (Ω = *|*Ω*|* in axes labels for brevity).

These sizes define the resolution with which the image is locally represented. The required/desired resolution is given everywhere by the *Implied Resolution Function*, which attributes high resolution to image areas where the intensity rapidly changes in space (e.g., edges), and low resolution to areas with low variation in intensity (e.g., background or uniform foreground). The Implied Resolution Function defines the radius of a neighborhood around each pixel. Within this neighborhood, the image intensity can be reconstructed at any location by taking a non-negative weighted average of the particles contained in it.

Determining optimal neighborhoods such that the reconstructed intensities are guaranteed to be closer to the original intensities than a user-defined threshold is the core of the problem. We here present an algorithm, called the *Pulling Scheme*, that efficiently solves this problem. The Pulling Scheme finds a set of particles, i.e. their locations and intensities, such that the required resolution is guaranteed everywhere and the image intensity can be reconstructed at each location from the intensities of the particles in its neighborhood with an accuracy that is guaranteed to be better than a user-defined threshold. Therefore, the APR efficiently finds a content-adaptive representation of the image with full user control over the representation quality.

If lossless representation is required, the APR places one particle at each pixel, in which case it becomes equivalent to the original pixel representation. However, fluorescence microscopy images are typically sparse, such that the number of particles can be orders of magnitude less than the number of pixels if small intensity deviations (e.g., within the imaging noise) are allowed. The computational and storage costs of the APR are proportional to the number of particles, and no longer to the number of pixels. By focusing on informative image areas, the APR reduces storage and computational costs and increases the information-to-data ratio.

## Reconstruction Condition

For the APR to optimally represent a given image with intensities *I*(*y*) at pixels *y*, the Implied Resolution Function should be as large as possible at every location, while still guaranteeing that the image can be reconstructed within the user-specified relative error *E* scaled by the Local Intensity Scale *σ*(*y*). The Local Intensity Scale *σ*(*y*) is an estimate of the range of intensities present locally in the image. Considering an arbitrary Resolution Function *R*(*y*), we can formulate the problem as finding the largest *R*(*y*) everywhere that satisfies

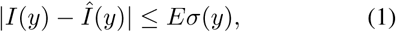

where 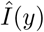 is the reconstructed intensity calculated by *any* non-negative weighted average over particles within *R*(*y*) distance of *y*. We call this the *Reconstruction Condition*. For the 1D example shown in Figure 2, a constant local intensity scale *σ*(*y*) = 1 is used. Maximizing *R*(*y*) minimizes 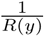, which is proportional to the locally required sampling density. Therefore, maximizing *R*(*y*) results in the minimum number of particles used. Unfortunately, finding the optimal *R*(*y*) that satisfies the Reconstruction Condition for arbitrary images requires a number of compute operations that proportional to the square of the number of pixels *N*. This computational cost is prohibitive even for modestly sized images. We propose two conservative restrictions on the problem and show that the optimal solution to the restricted problem can be computed with a total number of operations that is proportional to *N*.

## APR Solution

We outline the two problem restrictions, and how they are used to formulate an efficient linear-time algorithm for creating the APR.

### Resolution Bound

The first restriction on the Resolution Function *R*(*y*) requires that for all original pixel locations *y* it satisfies the inequality

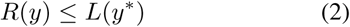

for all *y^∗^* with *|y − y^∗^| ≤ R*(*y*), and 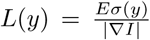. Here *|∇I|* is the magnitude of the image intensity gradient, which in 1D is 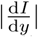 and can be computed directly from the image. We call this inequality the Resolution Bound, and *L*(*y*) the Local Resolution Estimate. If we assume the continuous intensity distribution underlying the image to be differentiable everywhere and the Local Intensity Scale *σ*(*y*) to be sufficiently smooth (See SuppMat 2.3), satisfying the Resolution Bound guarantees satisfying the Reconstruction Condition (See SuppMat 2.2). In Figure 2B, we illustrate that the Resolution Bound in 1D requires that a box centered at *y* of height *R*(*y*) and width 2*R*(*y*) does not intersect anywhere with the graph of *L*(*y*). Since the Resolution Bound represents a tighter bound than the Reconstruction Condition, the optimal solution to the Resolution Bound *R_b_* (*y*) is always less than or equal to the optimal solution to the Reconstruction Condition *R_c_*(*y*), therefore providing the same or a higher image representation accuracy. The dashed lines in Figure 2C illustrate this for the 1D example. As mentioned above, solving for the optimal Resolution Function requires computer time ∝ *N*^2^. However, we show next that the Resolution Bound can be found optimally with linear time in *N* if we restrict the Resolution Function to be composed of square blocks.

### Finding the Resolution Function with Particle cells

The second restriction is that the blocks constituting the Resolution Function must have edge lengths that are powers of 1/2 of the image edge length. The piecewise constant Resolution Function defined by the uppermost edges of these blocks is called the *Implied Resolution Function R^∗^*(*y*) and is shown in black in Figure 2C. The blocks we call *Particle Cells*. They have sides of length 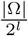, where |Ω| is the edge length of the image, measured in pixels. The number *l* is a positive integer we call the *Particle Cell Level*. Each Particle Cell *c_i,l_*, is therefore uniquely determined by its level *l* and location *i*. Figure 2C inset, illustrates these definitions for a single Particle Cell (See SuppMat 4 for the nD formal definition). The size of the blocks on the lowest resolution level is half the size of the image (*l*_min_ = 1), and the highest level of resolution *l*_max_ contains boxes the size of the original pixels. For image edge lengths that are not powers of 2, the parameter Ω is rounded upwards to the nearest power of two without padding the image.

Using these two restrictions, the problem of finding the optimal Resolution Function can be reduced to finding the smallest set *𝒱* of particle cells that defines an Implied Resolution Function *R^∗^*(*y*) that satisfies the Resolution Bound (SuppMat 4.1). We call this minimal set of Particle Cells the *Optimal Valid Particle Cell* (OVPC) set.

To construct an algorithm that efficiently finds the OVPC set for a given Local Resolution Estimate *L*(*y*), we first formulate the Resolution Bound in terms of Particle Cells. This formulation requires arranging the Particle Cells *c_i,l_* by level *l* and location *i* in a tree structure, as shown in Figure 2D. In 1D this is a binary tree, in 2D a quad-tree, and in 3D an oct-tree. When arranged as a tree structure, we can naturally define children and neighbor relationships between Particle Cells, as shown in green and blue in the example. We also define the descendants of a Particle Cell as the set of all children and children's children up to the maximum resolution level *l*_max_. Given these definitions, the Local Resolution Estimate *L*(*y*) can be represented as a set of Particle Cells *ℒ* by iterating over each pixel *y^∗^*, and adding the Particle Cell with level 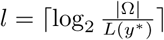 and location 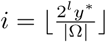 to *ℒ* if it is not already in *ℒ* (assuming the lower-left boundary of the image is at zero). Figure 2E illustrates how *ℒ* relates to *L*(*y*), with *ℒ* also represented in Figure 2D in the tree structure. We call this set of Particle Cells the *Local Particle Cell* (LPC) set *ℒ* (See SuppMat 4.2).

We can then represent the Resolution Bound in terms of *ℒ*. A set of Particle Cells *𝒱* will define an Implied Resolution Function that satisfies the Resolution Bound for *L*(*y*), if and only if the following statement is true: for every Particle Cell in *𝒱*, none of its descendants, or neighbors’ descendants, are in the LPC set L (SuppTheorem 1). We call any set of Particle Cells satisfying this statement valid. The OVPC set *𝒱* is then uniquely defined as the valid set for which replacing any combination of Particle Cells with larger Particle Cells would result in *𝒱* no longer being valid (SuppTheorem 2).

### Pulling Scheme

We present an efficient algorithm for finding the OVPC set *𝒱* called the *Pulling Scheme*. The name is motivated by how a single Particle Cell in *ℒ pulls* the resolution function down to enforce smaller Particle Cells across the image. The Pulling Scheme finds the OVPC set *𝒱* directly, without explicitly checking for validity or optimality. The result is by construction guaranteed to be valid and optimal. In order to derive the algorithm, we leverage three properties of OVPC sets:

1. *Predictable and self-similar structure*: Neighboring Particle Cells never differ by more than by one level and are arranged in a fixed pattern around the smallest Particle Cells in the set. This local structure is independent of absolute level *l* and endows the set with a self-similar structure. Using this structural feature, the OVPC set *𝒱* for a LPC set *ℒ* with only one Particle Cell *c_i,l_* can be generated directly for any *i* and *l*.
2. *Separability*: We can find the OVPC set given a LPC set *ℒ* by considering each cell in *ℒ* separately and then combining the smallest Particle Cells from all sets that cover the image (see Supp Lemma 1). SFigure 4 illustrates this separability property.
3. *Redundancy*: The redundancy property tells us that when constructing *𝒱*, we can ignore all Particle Cells in *ℒ* that have descendants in *ℒ*. This is because descendants provide equal or tighter constraints on the resolution function than their parent Particle Cells (see SuppLemma 2 for the proof).

These properties enable us to efficiently construct *𝒱* by propagating solutions from individual Particle Cells in *ℒ*, one level at a time, starting from the highest level (*l*_max_) of the smallest Particle Cells in *ℒ*. Here we use a simple implementation that explicitly represents all possible Particle Cells in an image pyramid structure^1^. The *Pulling Scheme* is summarized in Supplementary Algorithm 1, and SFigure 7 illustrates the steps for each level. SuppMat 5.5 and SuppMat 13.5 provide additional details. The computational cost of the algorithm scales with the number of Particle Cells in *𝒱*. Further, computing the OVPC set *𝒱* using the Pulling Scheme incurs a computational cost that is proportional to the number of pixels N for a fixed information-to-data ratio.

Using the Equivalence Optimization (See SuppMat 5.4 and SuppMat 5.7), the computational and memory costs of the Pulling Scheme can be further reduced by a factor of 2*^d^*, where *d* is the image dimensionality, while obtaining the same solution. A second optimization restricts the neighborhood of particle cells to further reduce the total number of particles used, as described in SuppMat 5.6. We use both optimizations for results presented in this paper. Ultimately, the only operation that needs to be computed on the full pixel image is a simple filtering operations for the gradient magnitude.

### Placing the Particles

*𝒫* Given the Implied Resolution Function computed by the Pulling Scheme, the last step of forming the APR is to determine the locations of the particles *𝒫*. Locations must be chosen so that around each pixel *y* there is at least one particle within a distance of *R*^*^(*y*). This requirement is easily satisfied by placing one particle at the center of each Particle Cell in *𝒱*. Specifically, for each Particle Cell *c_i,l_* in *𝒱*, we add a particle *p* to *𝒫* with location 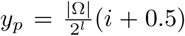. For each particle *p* we store the image intensity at that location *I_p_* = *I*(*y_p_*), interpolated from the original pixels as described in SuppMat 6. This way of arranging the particles has the advantage that the particle positions do not need to be explicitly stored, as they are determined by *𝒱*.

### Forming the APR= {*𝒱*, *𝒫*}

In Figure 3 we outline the steps required to form the APR from an input image. The APR can be stored as the combination of {*𝒱*, *𝒫*}. We represent the OVPC set *𝒱* by storing the integer level *l* and the integer location *i* for each Particle Cell. *𝒱* therefore defines the Implied Resolution Function *R^∗^*(*y*) for all *y* in the image. *𝒫* stores the intensities of all particles.

**Figure 3.**
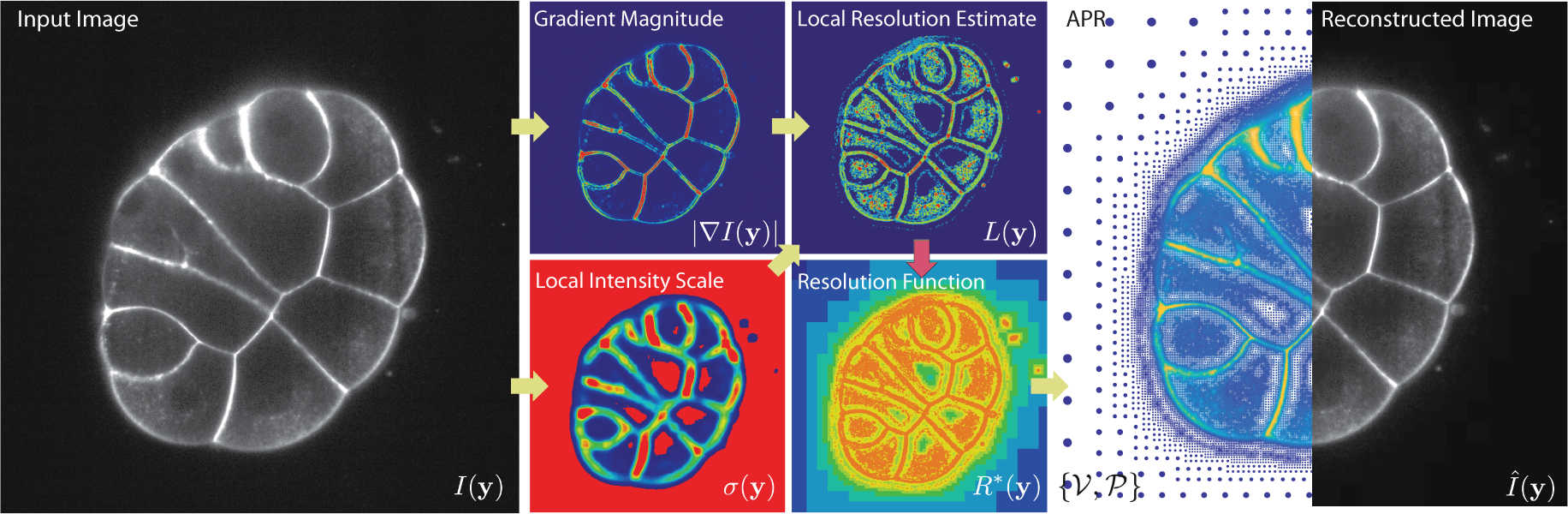
Procedure for forming the APR in 3D. The illustration shows a 2D slice of a fluorescence image (*Input Image I*(**y**), Dataset 10 in STable 3, courtesy of Lemaire lab, CRBM (CNRS) and Hufnagel lab, EMBL). First, the Local Intensity Scale *σ*(**y**) and the intensity Gradient Magnitude |∇*I*(**y**)| are calculated from the input image. These two are then combined to compute the Local Resolution Estimate *L*(**y**). The Pulling Scheme (*red arrow*) then uses *L*(**y**) to compute the optimal Implied Resolution Function *R^∗^*(**y**). This is then used to define the Optimal Valid Particle Cell set *𝒱* and the particle locations *𝒫*, which together form the APR (*right*). The left half of the *right* panel shows the particles of the APR with color encoding intensity and size encoding local resolution. The right half shows a piecewise constant reconstruction 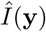 of the image for visualization.

### Imaging noise and accuracy

Determining *L*(*y*) requires computing the intensity gradient ∇*I* over the input image. In practice, the pixel intensities are noisy, which leads to uncertainty in the computed *L*(*y*). In SuppMat 7, we provide theoretical results how this uncertainty imposes a lower bound on the achievable representation accuracy *E*.

## 3D Fluorescence APR Implementation

We assess the properties of the APR for noisy 3D fluorescence microscopy images. Figure 3 illustrates the main steps of the implementation using a 2D image slice of a 3D image.

When implementing the APR, three design choices have to be made: First, one has to decide how to calculate the gradient magnitude |∇*I*(**y**)|. Second, one has to decide how to compute the Local Intensity Scale *σ*(**y**). Third, one has to decide how to interpolate the image intensity at particle locations *Ip* = *I*(**y***p*). Full details are given in SuppMat 13.

To calculate the gradient magnitude over the input image we use smoothing cubic B-Splines (*27*), which provide robust gradient estimation in the presence of noise. They require the setting a smoothing parameter *λ* depending on the noise level, as described in SuppMat 13.

For the Local Intensity Scale *σ*(**y**), we use a smooth estimate of the local dynamic range of the image, as described in SuppMat 13.3. This form of the local intensity scale accounts for variations in the intensities of labeled objects, similar to gain control in the human visual systems. We ensure that *σ* is sufficiently smooth (see SuppMat 4.4) by computing it over the image down-sampled by a factor of two. Examples are shown in Figures 1B and 3. The size of the smoothing window is given by a rough estimate of the standard deviation of the point-spread function (PSF) of the microscope. Further, a minimum threshold is introduced to prevent resolving background noise (see SuppMat 13).

Two methods are combined to interpolate pixel intensities to particle locations: for particles in Particle Cells at pixel resolution, the intensities are directly copied from the respective pixels, while for particles in larger particle cells, we assign the average intensity of all pixels in that Particle Cell (*19*).

We also provide a method for reconstructing a pixel image 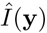 from the APR. A pixel image satisfying the Reconstruction Condition in Eq. 1 can be reconstructed from the APR using any non-negative weighted average of particles within *R^∗^*(*y*) of pixel *y*. In SuppMat 10 we discuss possible weight choices, providing examples of smooth, piecewise constant, and worst-case reconstructions. For displaying figures and benchmarking, unless otherwise stated, we use the piecewise constant reconstruction in this paper. This reconstruction sets all pixels inside a Particle Cell equal to the intensity of the particle in that cell and thus has the best computational efficiency.

All design decisions have been made to optimize robustness against imaging noise and computational efficiency. We find that the method is stable with respect to the choice of parameters. A discussion of parameter selection for real datasets is given in SuppMat 14, and the parameter values used for our test datasets are given in STable 3.

### Data structures

Appropriate data structures must be used to store and process on the APR efficiently. Ideally, these structures allow direct memory access at low overhead. Here, we propose a multi-level data structure for the APR, as described in SuppMat 18. Each APR level *l* is encoded similar to sparse matrix schemes with Particle Cell locations {*i_x_, i_y_, i_z_*}. This data structure efficiently stores *𝒱* and *𝒫* by explicitly encoding only one spatial coordinate (*i_y_*) per Particle Cell, while allowing random access. We call this data structure the *Sparse APR* (SA) data structure. It relies on storing one red-black tree of Particle Cell locations *i_y_* for each combination of {*i_x_, i_z_, l*}, caching access information for contiguous blocks of Particle Cells. When storing image intensity using 16 bits, the SA data structure requires approximately 50% more memory than the intensities alone. Simpler data structures, without the red-black tree, can be used to reduce this overhead if random access is not required. In all results presented here, we use the SA data structure.

### APR image file format

We store the APR using the HDF5 file format (*28*) and the BLOSC HDF5 plugin (*29*) for lossless Zstd compression of the Particle Cell and intensity data in the file.

## Validation

All benchmarks use the open-source C++ APR software library *LibAPR* (github.com/cheesema/LibAPR) compiled with with gcc 5.4.0 and OpenMP shared-memory parallelism on a 10-core Intel Xeon E5-2660 v3 (25 MB cache, 2.60 GHz, 64 GB RAM) running Ubuntu Linux 16.04. SuppMat 16 provides a detailed description of each benchmark and the parameters used.

### Benchmarks on synthetic data

We first assess the performance of the APR using synthetic benchmark data. SuppMat 15 and SFigure 26 outline the synthetic data generation pipeline. The key advantage of synthetic data is that all relevant image parameters can be varied and the ground-truth image is known. Synthetic images are generated by placing a number of blurred objects into the image domain and corrupting with modulatory Poisson noise. We study the influence of image size, content, and noise level on the performance of the APR. Spherical objects are used for simplicity unless otherwise indicated.

#### Reconstruction Condition

We experimentally confirm that the APR satisfies the Reconstruction Condition in Eq. 1 in the absence of noise. Figure 4A shows the empirical relative error 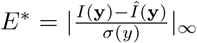 for increasing imposed error bounds *E*. In all cases, *E*^*^ < *E*, as required by the Reconstruction Condition. As expected, the number of particles used by the APR to represent the image decreases with increasing *E* (right axis). The results are unchanged when using more complex objects than spheres or different reconstruction methods (SFigure 29). Figure 4C provides examples of the quality of APR reconstruction at different *E*, compared to ground truth. In the absence of noise, the APR satisfies the Reconstruction Condition everywhere, guaranteeing a reconstruction error below the user-specified threshold, fulfilling the first part of RC1.

**Figure 4.**
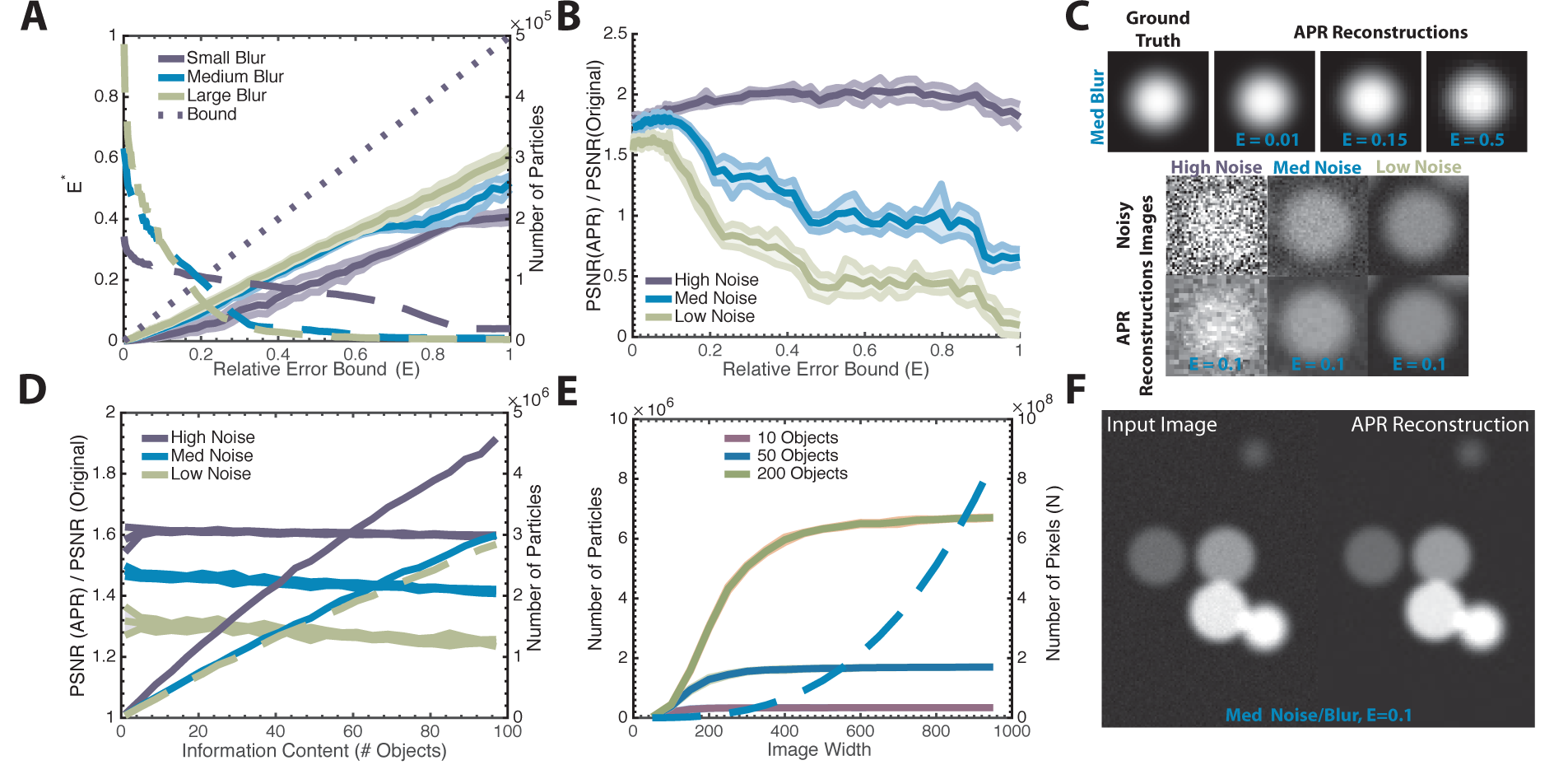
Benchmarking the APR on synthetic data. All results are shown with mean (*lines*) and standard deviation (*bands*). **A.** Observed reconstruction error *E^∗^* (solid lines, *left axis*) between the ground truth and the piecewise constant APR reconstruction (SuppMat 16.2) for noise-free images. Number of particles used by the APR (dashed lines, *right axis*) for different user-defined error bounds *E* (original image *N* = 128^3^ ≈ 2.1 × 10^6^). Results are shown for images of different sharpness (blur) (*inset legend*). The APR reconstruction error is below the specified bound in all cases (dotted line). More accurate APRs require more particles. **B.** Peak signal-to-noise ratio (PSNR) of the APR relative to the PSNR of the original pixel image for different error bounds *E* and image noise levels (*inset legend*) (SuppMat 16.3). For low *E*, the APR has a better PSNR than the input images. **C.** Examples of test images of spherical objects with different noise levels and *E* used in the benchmarks. The *top row* shows the APR reconstructions of the medium-blur noise-free test image at different *E* compared to the ground truth. The *bottom rows* compare the original noisy images with their APR reconstructions for *E* = 0.1, illustrating the inherent denoising property of the APR. **D.** PSNR ratio (solid lines, *left axis*) and number of particles used (dashed lines, *right axis*) for images containing different numbers of objects, i.e., different information content, for *E* = 0.1. (SuppMat 16.4). In all cases, the PSNR of the APR is better than that of the input image, and the number of particles scales at most linearly with image information content (original image *N* = 300^3^ = 2.7 × 10^7^). **E.** Number of APR particles (solid lines, *left axis*) and input image pixels (dashed lines, *right axis*) for images of different image sizes containing a fixed number of objects (SuppMat 16.5) (E = 0.1). The number of particles plateaus as soon as the objects in the image are well resolved. **F.** Visual comparison of a medium-blur, medium-noise image containing six objects (*left*) with its piecewise constant APR reconstruction (*right*) for *E* = 0.1.

#### Robustness against noise

In real applications, images are corrupted by noise. We find that the introduction of noise introduces a lower limit on the error *E^∗^* that can be achieved (see first plot in SFigure 30). This observation agrees with theoretical analysis (SuppMat 7.3). This lower bound is entirely due to the noise in the pixel intensity values, while the adaptation of the Implied Resolution Function *R^∗^*(*y*) is robust to noise. This is demonstrated in the second plot in SFigure 30, where noisy particle intensities are replaced with ground-truth values for the reconstruction step. Adaptation is still done on the noisy pixel data. Again, *E^∗^* can be made arbitrarily small, indicating that the construction of the APR is robust against imaging noise. This result also agrees with the theoretical analysis of the impact of errors in *L*(*y*) on the Implied Resolution Function (SuppMat 7.2).

To understand how to best set *E* in the presence of noise, we compute the observed peak signal-to-noise ratio (PSNR) of the reconstructed image and compare with the PSNR of the original image. Figure 4B shows that decreasing *E* to zero does not maximize the PSNR. Instead, for medium to high quality input images, the PSNR is highest between an *E* of 0.08 and 0.15. For low-quality input images, we find a monotonic relationship between the PSNR and *E*, as de-noising from downsampling dominates. Also, for *E* < 0:2 the reconstruction error is always less than the noise in the input image, reflected in a PSNR ratio greater than one. Therefore, for noisy images with medium to high quality, there is an optimal range for *E* between 0.08 and 0.15. In this range, the reconstruction errors are less than the imaging noise, and the signal-to-noise ratio of the APR is better than that of the input pixel image, fulfilling also the second part of RC1.

The noise distribution over the particles in the APR depends on the original noise distribution of the pixel image and on the method used to interpolate the particle intensities from the pixels. In SuppMat 7.6, we provide both numerical and theoretical results for the interpolation scheme used here. We consider both Gaussian and Poisson noise on the input pixel image. The variance of the noise scales inversely proportional to the Particle Cell level *l*. For Gaussian noise, the noise remains Gaussian on each level with variance scaled by a factor of 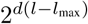, where *d* is the image dimension. This is expected, as coarser levels correspond to more averaging and hence noise reduction.

#### Response to image content

In Figure 4D we show how the APR adapts to image content. This adaptation is manifested in the linear relationship between the number of objects (spheres) randomly placed in the image and the number of particles used by the APR (*right axis*). Adaption is linear despite the brightness of the objects randomly varying over an order of magnitude (see SuppMat 16.4). Image quality is maintained throughout (*left axis*). Figure 4C shows an example of a medium-quality input image and its APR reconstruction. Figure 4E shows that the number of particles used by the APR to represent a fixed number of objects becomes independent of image size. Also, if pixel resolution and image size are increased proportionally, the APR approaches a constant number of particles (S Figure 31). These results show that the APR adapts proportional to image content, independent of the number of pixels, fulfilling RC2.

#### Local Intensity Scale

So far, we have not directly assessed the validity of the Local Intensity Scale *σ*. In order to do this, we need a ground-truth reference. In SuppMat 15.5 we introduce the *perfect APR*, and the *Ideal Local Intensity Scale σ*^ideal^ that can be calculated for synthetic data. This ground-truth representation is then used to benchmark the APR.

The results in STables 1 and 2 show that the local intensity scale we use is effective over wide range of scenarios. However, for crowded images with large contrast variations (two orders of magnitude or more), we find that the Local Intensity Scale over-estimates the dynamic range of dim regions that are close to bright regions. This effect is most pronounced in high-quality images, where alternative formulations of the Local Intensity Scale could provide better results.

#### Computational cost

Due to the adaptivity of the APR, its computational cost depends on image content through the number of particles, and not on the input image size *N*. For a given input image, we define the *Computational Ratio* (CR) as:

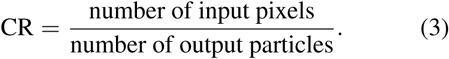

We assess the performance of the APR for synthetic images with numbers of objects roughly corresponding to CR = 5, 20, 100, representing high, medium, and low complexity images (SFigure 32, SuppMat 17.1). The results are given in Table 1. The APR achieved effective CR values of 5.63, 19.7, and 93.9, respectively.

**Table 1.** Summary statistics of the APR benchmarks on synthetic and real-world images. Results are shown for synthetic images with fixed CR = 5,20,100 and for 19 real-world exemplar datasets (see STable 3). For the exemplars, we report the means, standard deviations (brackets), and medians of the values over all exemplar images. For the synthetic fixed-CR benchmarks, the effective CR and the Memory Compression Ratios (MCR) are averaged over image sizes from 200^3^ to 800^3^ (standard deviations in brackets) and the values for absolute runtimes and storage requirements are given for images of size 800^3^. For comparison, we also report the MCR using lossy *within-noise-level* (WNL) compression (*30*) of both the APR and the pixel images for the same compression parameter value (*q* = 2, see SuppMat 20.1). We also show the time taken to transform the images to the APR on the benchmark machine and the runtime of the Pulling Scheme alone.

| | Computational<br>Ratio (CR) | Raw Image<br>Size (GB) | Compressed<br>APR GB | MCR of APR | MCR of APR-WNL, $q=2$ | MCR of<br>Pixels-WNL,<br>$q=2$ | Pipeline<br>Time (s) | Pulling Scheme<br>Runtime (s) |
| --- | --- | --- | --- | --- | --- | --- | --- | --- |
| *CR5* | 5.63 (0.02) | 1.024 | 0.129 (0.0006) | 7.9 (0.04) | 19.6 (0.97) | 5.4 (0.36) | 2.34 (0.086) | 0.104 (0.002) |
| *CR20* | 19.7 (0.13) | 1.024 | 0.036 (0.0002) | 28.4 (0.19) | 64.4 (2.03) | 5.69 (0.62) | 2.01 (0.07) | 0.04 (0.003) |
| *CR100* | 93.9 (1.6) | 1.024 | 0.007 (0.0001) | 139.9 (2.1) | 282 (66.4) | 5.85 (0.75) | 1.87 (0.08) | 0.027 (0.005) |
| *Exemplars Mean* | 51.1 (89.3) | 1.869 (1.38) | 0.051 (0.053) | 129.5 (284) | 297.8 (593) | 95.8 (166) | 3.65 (2.19) | 0.10 (0.08) |
| *Exemplars Median* | 22.7 | 1.258 | 0.027 | 36.8 | 107.1 | 33.1 | 2.19 | 0.066 |

### Benchmarks on real data

We present results for a corpus of 19 exemplar volumetric fluorescence microscopy datasets of different content and imaging modalities, ranging in size from 160 MB to 4 GB. The datasets and parameters used are described in STable 3&4 and summary statistics in Table 1. SFigure 33 shows a cross-section of the APR for exemplar dataset 7, and SVideo 1 illustrates the Implied Resolution Function and APR reconstruction for exemplar dataset 1. A comparison of the APR with Haar wavelet thresholding for natural scene images (*31*) is given in SuppMat 12.

#### Memory requirements

Calculation of the APR from an image requires approximately 2.7 times (for 16-bit images) the size of the original image in memory. The maximum image size is only limited by available main memory (RAM) of the machine and by the ability to globally index the particles using an unsigned 64-bit integer. Our pipeline has been successfully tested on datasets exceeding 100 GB (SFigure 35). To test on very large data, exemplar dataset 17 was tiled 200 times to create a 320 GB image. Using the same parameters as for the original image resulted in an APR of 4.08 GB and hence CR of 20.2, compared to a CR of 21 for the original image.

#### Execution time

On our benchmark system, we find linear scaling in *N* and an average data rate of 507 MB/s for transforming images to their APR. This rate corresponds to 3.9 seconds to form the APR from an input image of size *N* = 1000^3^. On the exemplars, execution times range from 0.37 seconds to 8.14 seconds, with an average of 3.65 seconds. Table 1 summarizes the results. We find the following distribution of computation time: the Pulling Scheme on average takes less than 3.5% of the total time while the computation of the gradient magnitude using smoothing B-splines dominates the execution time, taking up to 59% of the total time. For details see SuppMat 19.

The pipeline shows efficient parallel scaling (Amdahl's Law, parallel fraction = 0.95) on up to 47 cores, achieving data rates of up to 1400 MB/second (SFigure 35). This enables real-time conversion of images to the APR, as it is faster than the acquisition rate of microscopes (*32, 33*).

We conclude that images can be rapidly converted into the APR with a cost that scales at most linearly with image size *N*, fulfilling RC3.

#### Storage requirements

For the fixed-CR datasets, we observe an average Memory Compression Ratio^2^ of 1.4 times the CR. The median MCR of the exemplars is 36.8, and the mean is 129.5. This corresponds to an average size of the input images of 1.87 GB and 51 MB on average for the compressed APR files. Table 1 summarizes the results and STable 4 provides the image details.

When the APR is stored on disk, on average 89% of the bytes are used to store the particle intensities, implying that the APR data structures occupy 11% on average. In addition, the APR particle intensities can be compressed further in a lossy manner using existing lossy image compression algorithms. This is shown in Table 1, where we also report the MCR using the *within-noise-level* (WNL) compression algorithm for large fluorescence images (*30*) for both the APR and original pixel image. Details on the implementation and benchmarks on synthetic data are provided in SuppMat 20.1. On synthetic data, we find the APR and pixel image provide the same image quality upon lossy compression, but the APR increases the compression ratio five fold. This indicates that the APR data structures are better suited for further compression using existing lossy compression techniqies.

In summary, the APR can be efficiently compressed with a file size proportional to the image content, fulfilling RC2. Unlike compression techniques, the APR is an image *representation* that can be leveraged to accelerate downstream processing tasks, including compression, without reverting to the original pixel image.

## Image Processing on the APR

We show how the APR reduces the memory and computational cost of downstream image-processing tasks (RC4). Once we have transformed the input image into an APR, the input image is no longer needed. All processing, storage, and visualization can be done directly on the APR.

Image-processing methods are always developed using a certain interpretation of images. Just like pixels, one can also interpret and use the APR in different ways depending on the processing task. These interpretations align with those commonly used in pixel-based processing. Figure 5A-D outlines the four main interpretations of the APR: collocation points, continuous function approximations, trees, and graphs.

**Figure 5.**
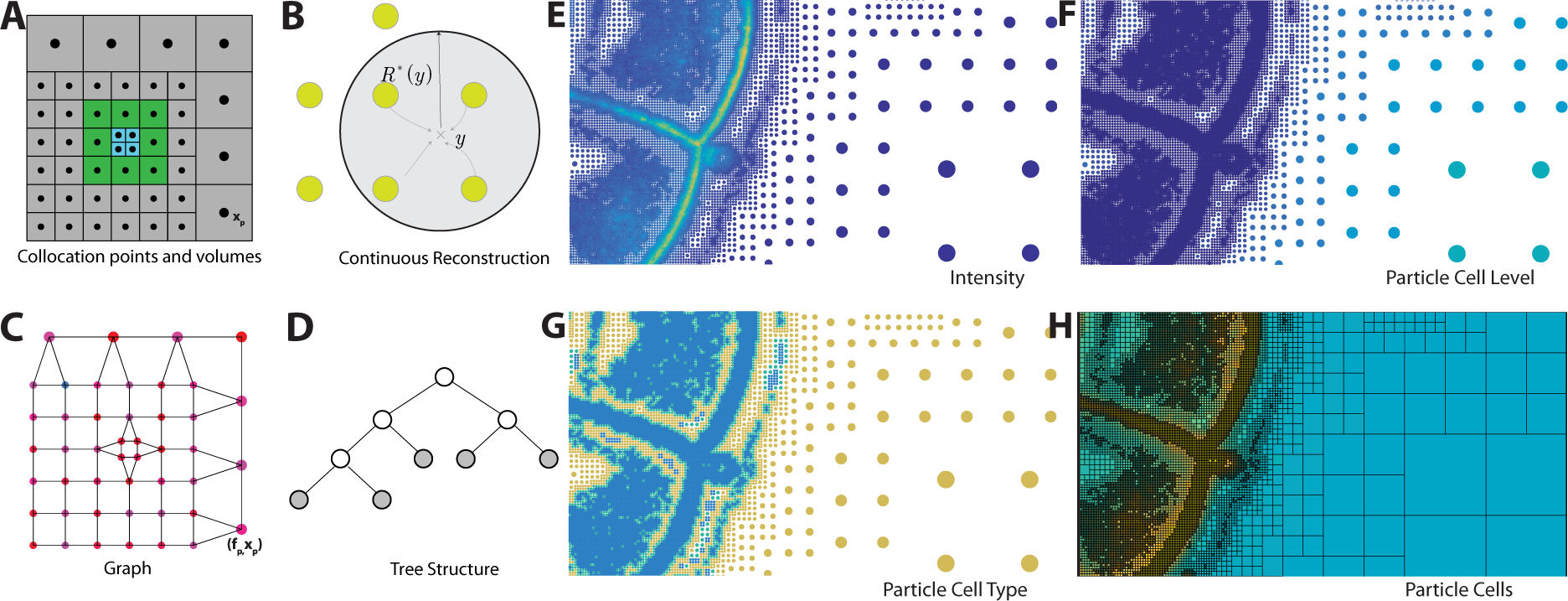
Interpretations of the APR for image processing. **A.** The APR can be interpreted as a spatial partition defined by the Particle Cells in *𝒱*, or by the set of particles *𝒫* with positions *x_p_*. This interpretation relates to the concept of super-pixels (*13*). **B.** The APR can be interpreted as a continuous function approximation where the intensity value can be reconstructed at each location *y*, also between particles and pixels, relating to smooth particle function approximations (*34*). **C.** The APR can be interpreted as a graph, where the particles are vertices and edges link neighboring particles (SuppMat 21). This relates the APR to graphical models of pixel images (*35*). **D.** The APR can be interpreted as a pruned binary tree (quad-tree in 2D, oct-tree in 3D) with links between parent and children Particle Cells. This relates the APR to wavelet decompositions (*17*), image pyramids (*19*), and tree-based methods (*36*). **E**–**H.** While particles store local fluorescence intensity, just like pixels (**E**), they also provide additional information that is not available on the pixels. This includes the Particle Cell level containing information about the local level of detail in the image (**F**), the Particle Cell type encoding the structure of the image (**G**), and the Particle Cell membership providing a content-adaptive disjoint partition of the image domain (**H**).

### Performance metrics

The APR can accelerate existing algorithms in two ways: First, by decreasing the total processing time through reducing the number of operations that have to be executed. Second, by reducing the amount of memory required to run the algorithm. The relative importance of the two, and the degree of reduction, depends on the specific algorithm and its implementation. We use quantitative metrics to evaluate the improvements for different algorithms and input images.

The first evaluation metric relates to the computational performance of the algorithm. For a given algorithm and implementation, we define the speed-up (SU) as:

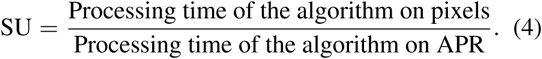

It is insightful to relate the SU to the CR by SU = CR * (Pixel-Particle Speed Ratio) (PP), where PP = (Time to compute the operation on *one* pixel)/(Time to compute the operation on *one* particle). The value of PP depends on many factors, including memory access patterns, data structures, hardware, and the absolute size of the data in memory. Consequently, even for a given algorithm running on defined hardware, the PP is a function of the input image size *N*. For tasks with PP *<* 1, as in some low-level vision tasks, there is a minimum value of CR above which the algorithm is faster on the APR than on pixels. For tasks with PP > 1, processing on the APR is always faster than on pixels.

For an algorithm run on a pixel image, in most cases the Memory Cost (MC) in bytes scales linearly with the number of pixels *N* and the number of required temporary and output variables (i.e. copies of the image), as MC=(Number of variables)*(Data type in bytes)\**N*. The memory cost of the APR

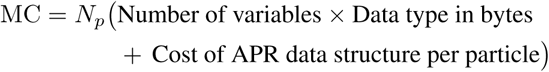

where *N_p_* is the number of particles, and the cost of the data structure per particle depends on *N*. We find an estimated average of 8 bits per particles overhead for the Sparse APR data structure. As the number of algorithm variables increases, the overhead of the APR is amortized so that the reduction in memory cost approaches the CR.

### Image Processing Performance Bench-marks

We analyze two low-level and one high-level image-processing tasks. These are neighbor access and filtering as low-level tasks, and image segmentation as high-level task. The low-level tasks represent a lower bound on the benefits of the APR due to their simple operations and access patterns, which are best suited for processing on pixels. The segmentation task in contrast provides a representative practical example of microscopy image analysis.

For these three benchmarks, we provide results for the computational and memory metrics for three fixed-CR datasets with input images from *N* = 200^3^ up to *N* = 1000^3^, and for all real-world exemplar datasets. The results of all benchmarks are summarized in Table 2. SuppMat 22 describes the benchmark protocols.

**Table 2.** Summary statistics of the image-processing benchmarks on synthetic and real-world images. For the exemplars, we report the means (standard deviation in brackets) of the values over all exemplar images. For the synthetic fixed-CR datasets, the speed-ups (SU), Pixel-Particle Speed Ratios (PP), and Memory Reduction Ratios (MRR) = (Memory Cost Pixels)/(Memory Cost APR) are averaged over image sizes from 200^3^ to 1000^3^; absolute timings and memory requirements are given for images of size 800^3^. Graph-cut segmentation on pixels was not possible for 800^3^ images, as the memory requirement exceeded the 64 GB available on the benchmark machine. The corresponding entries in the table (marked with *^∗^*) are extrapolated from benchmarks run on smaller images and the SU, PP, and pixel timing for the exemplars could not be determined in this case (*N/A*). See main text and SuppMat 22 for a detailed descriptions of the benchmarks.

|  | Speed Up | Time APR (s) | Time Pixels (s) | PP | Memory Pixels (GB) | Memory APR (GB) | MRR |
| --- | --- | --- | --- | --- | --- | --- | --- |
| **Linear Neighbor Iteration** |  |  |  |  |  |  |  |
| *CR5* | 0.55 (0.02) | 1.86 (0.07) | 1.02 (0.0002) | 0.097 (0.003) | 3.072 (0) | 0.599 (2.7) | 5.12 (0.02) |
| *CR20* | 1.9 (0.09) | 0.54 (0.03) | 1.02 (0.002) | 0.096 (0.004) | 3.072 (0) | 0.181 (1.0) | 16.9 (0.09) |
| *CR100* | 7.1 (0.5) | 0.14 (0.009) | 1.02 (0.007) | 0.076 (0.005) | 3.072 (0) | 0.053 (0.0005) | 60.2 (2.3) |
| *Exemplars Mean* | 4.06 (5.7) | 0.86 (0.5) | 1.83 (1.3) | 0.094 (0.01) | 5.61 (4.2) | 0.278 (0.28) | 37.5 (56) |
| **Random Neighbor Access** |  |  |  |  |  |  |  |
| *CR5* | 0.71 (0.03) | 15.4 (0.2) | 11.0 (0.4) | 0.126 (0.005) | 3.072 (0) | 0.599 (2.7) | 5.12 (0.02) |
| *CR20* | 3.52 (0.3) | 3.23 (0.05) | 11.4 (0.8) | 0.178 (0.01) | 3.072 (0) | 0.181 (1.0) | 16.9 (0.09) |
| *CR100* | 24.8 (0.8) | 0.44 (0.01) | 11.01 (0.3) | 0.26 (0.007) | 3.072 (0) | 0.053 (0.0005) | 60.2 (2.3) |
| *Exemplars Mean* | 11.57 (23.6) | 7.29 (10.1) | 21.4 (16) | 0.17 (0.05) | 5.61 (4.2) | 0.278 (0.28) | 37.5 (56) |
| **Image Filtering** |  |  |  |  |  |  |  |
| *CR5* | 7.36 (1.2) | 1.36 (0.009) | 8.02 (0.2) | 1.26 (0.2) | 4.10 (0) | 0.93 (0.003) | 4.38 (0.04) |
| *CR20* | 14.82 (3.7) | 0.76 (0.01) | 8.07 (0.3) | 0.77 (0.2) | 4.10 (0) | 0.30 (0.002) | 12.82 (0.9) |
| *CR100* | 31.10 (13) | 0.57 (0.003) | 7.96 (0.3) | 0.35 (0.15) | 4.10 (0) | 0.09 (0.0002) | 36.85 (6.7) |
| *Exemplars Mean* | 12.27 (3.0) | 1.24 (0.93) | 14.13 (9.8) | 0.51 (0.33) | 7.48 (5.5) | 0.36 (0.28) | 24.49 (19) |
| **Image Segmentation** |  |  |  |  |  |  |  |
| *CR5* | 5.10 (0.7) | 1.87 (0.02) | 8.86 (0.09) | 0.86 (0.04) | $\approx 68.5^*$ (0) | 12.57 (0.08) | 5.51 (0.14) |
| *CR20* | 18.30 (2.6) | 0.48 (0.003) | 8.83 (0.08) | 0.95 (0.07) | $\approx 68.5^*$ (0) | 3.75 (0.02) | 18.18 (0.3) |
| *CR100* | 85.3 (12) | 0.10 (0.001) | 8.78 (0.09) | 0.97 (0.06) | $\approx 68.5^*$ (0) | 0.80 (0.003) | 84.09 (2.9) |
| *Exemplars Mean* | N/A | 6.99 (5.9) | N/A | N/A | $\approx 385^*$ (286) | 13.54 (11.7) | 39.72 (40) |

#### Neighbor Access

For each pixel or particle, the task involves averaging the intensities of all face-connected neighbors (see SuppMat 22.1 for details). In the APR, neighbors are defined by the particle graph, as shown in Figure 5C and described in SuppMat 21. We bench-mark two forms of neighbor access: *Linear iteration* loops over all neighbors in sequential order. *Random access* visits neighbors in random order, irrespective of how they are stored in memory.

For linear iteration, the APR shows low speed-ups. It is even slower than pixel operations for images with CR = 5 and for four of the exemplar datasets (Table 2, group 1). This is because linear iteration is optimally suited to pixel images. However, the APR provides consistently higher speed-ups for random neighbor access, especially for high CRs. This is likely due to the smaller overall size of the APR improving cache efficiency.

The total memory cost of the APR reflects the CR of the dataset. This provides significant memory cost reductions across all benchmark datasets for both the linear and random neighbor access patterns.

#### Image filtering

We consider the task of filtering the image with a Gaussian blur kernel (see SuppMat 22.2). We exploit the separability of the kernel and perform three consecutive filtering steps using 1D filters in each direction. On the APR, this requires locally evaluating the function reconstruction. The benchmark results are shown in Table 2, group 3. Directly filtering the APR consistently outperforms the pixel-based pipeline, both in terms of memory cost and execution time.

In SuppMat 22.2.2 we analyze the results in detail and find that the APR is most appropriate if the filtering result has a similar structure as the original image, such that the same set of content-adapted particles is suitable to represent both images. SFigure 40 illustrates this, showing that for a weak blur the APR filter has higher PSNR than the pixel filter. For stronger blurs this is reversed, because the specific APR adapted to the input image is no longer optimal to represent the filtered image.

#### Image segmentation

We perform binary image segmentation using graph cuts, using the method and implementation of Ref. (*35*) to compute the optimal fore-ground/background segmentation for both APR and pixel images. When computing the cut energies, we exploit the additional information provided by the particle cell level, type, and local intensity range. To allow direct comparison with pixel-based segmentation, we interpolate all energies calculated on the APR to pixels and determine the cuts over the pixel image using the same energies. For both APR and pixel images, a face-connected neighborhood graph is used. Given the energy calculations are identical, we benchmark the execution time and memory cost of the graph-cut solver. The results are shown in Table 2, group 4. For the APR we find speed-ups directly reflecting the CR.

Using the APR, all examplar images can be segmented without problems, illustrating the benefits of the reduced memory cost of the APR, while pixel images can only be segmented for sizes *N* ≤ 550^3^ on our benchmark machine with 64 GB RAM. Image-analysis algorithms with high memory overhead, such as graph-cuts, thus benefit from the APR particularly strongly.

We validate the APR segmentations by comparing both the APR and pixel-based segmentations to ground truth using the Dice coefficient (*38*). Across datasets, we find that the Dice coefficients are not statistically significantly different (p-value: 0.92, Welch's t-test). We provide a representative example in SVideo 2 and show a 3D rendering of a segmentation in Figure 6E. For more details, see SuppMat 22.3.

**Figure 6.**
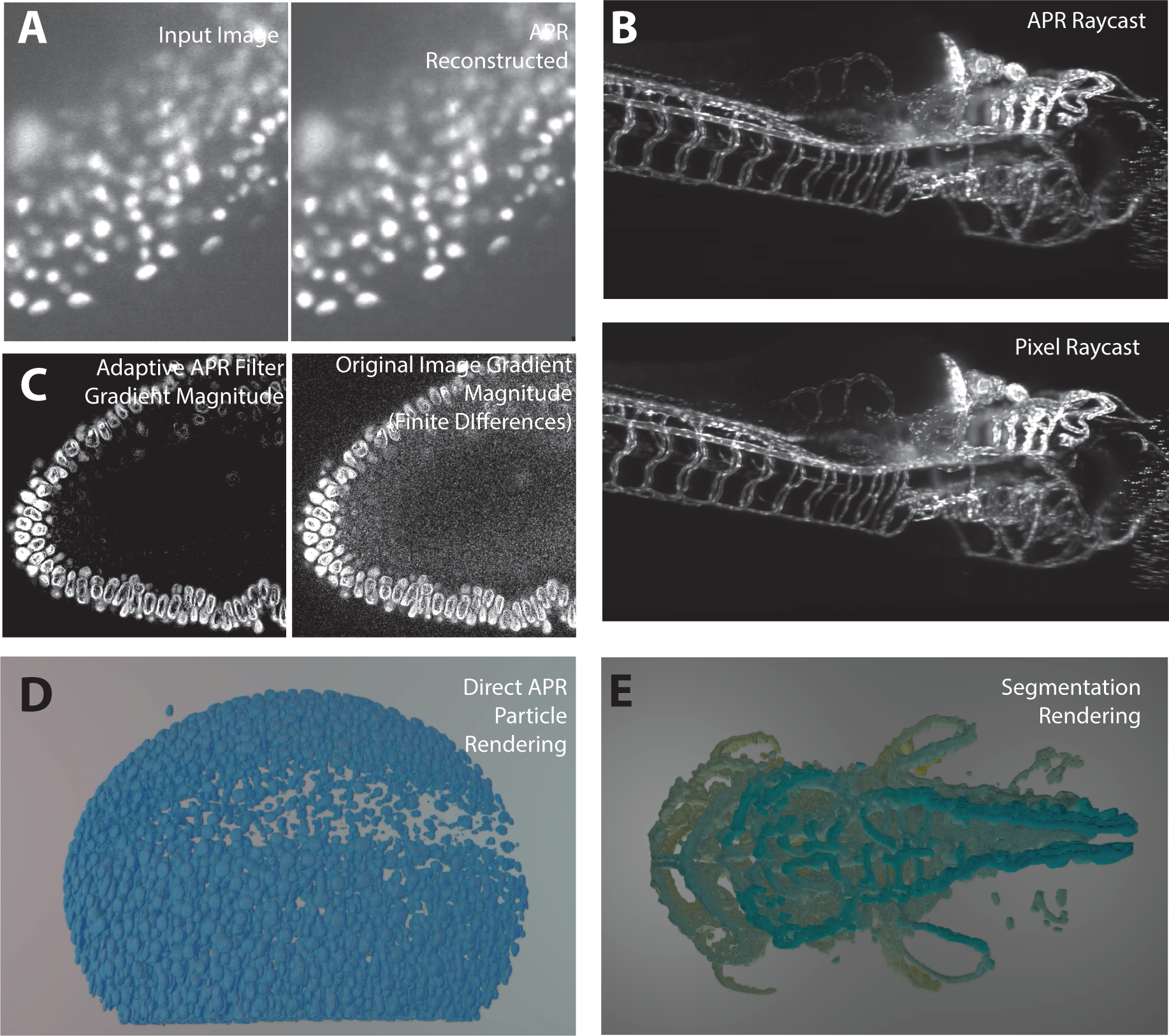
Image processing using the APR. **A.** Comparison of an example image (*left*, exemplar dataset 7) with its piecewise constant APR reconstruction (*right*). **B.** Comparison of the maximum-intensity projection of a direct 3D APR raycast (*top*) with the maximum-intensity projection of the pixels (*bottom*) for exemplar dataset 17 (full image see SFigure 45). **C.** Comparison of the intensity-gradient magnitude estimated using the Adaptive APR Filter (*left*, SuppMat 22.4) and central finite differences over the pixels (*right*) for exemplar dataset 6 (Tomancak Lab, MPI-CBG). **D.** Direct 3D particle rendering of Zebrafish nuclei (exemplar dataset 7) using the open-source visualization tool *Scenery* (*37*). **E.** APR Volume rendering of a 3D image-segmentation result, colored by depth, computed using graph-cut segmentation directly on the APR, as described in SuppMat 22.3.3 (exemplar dataset 13, cf. **B**). Segmentation on the APR took 5.5 seconds. Segmentation on the pixel image was impossible on our benchmark machine due to memory requirements. (Raw images in (A,B,D,E) courtesy of Huisken Lab, MPI-CBG & Morgridge Institute for Research.)

### Novel Algorithms

The APR provides additional information about the image that is not contained in pixel representations. This information can be exploited in image-processing algorithms, as illustrated in the segmentation example above. In addition, it can also be used to design entirely novel, APR-specific algorithms, as demonstrated in the following example.

#### Adaptive APR filter

We define a discrete filter over neighboring particles in the APR particle graph. Since the distances between neighboring particles vary across the image, depending on image content, this amounts to spatially adaptive filtering with the filter size automatically adjusting to the content of the image. On the APR, this only requires linear neighbor iteration, while an adaptive pixel implementation would be significantly more complex.

SuppMat 22.4 describes the adaptive APR filter in detail. SFigure 42 shows synthetic results for an adaptive blurring filter, and SFigure 43 for a filter that adaptively estimates the intensity gradient magnitude. In both examples, the adaptive APR-filtered results have higher PSNR than results from corresponding non-adaptive pixel filters, also demonstrated in Figure 6C.

### Visualization

Images represented using the APR can directly be visualized without going back to pixels. The APR image can be visualized using both traditional and novel visualization methods. We provide examples of the following visualization methods:

*Visualization by slice*: Figure 6A and SVideo 1 show examples of a slice-wise APR reconstruction in comparison with the pixel images.

*Raycasting* for direct 3D visualization of an APR. Figure 6B and SVideo 3 show a perspective maximum-intensity projection in comparison with the same ray-cast of the original pixel image. The resulting visualizations are visually indistinguishable. APR raycasting only requires storing and computing on the APR, reducing memory and computational costs proportionally to the CR of the image, enabling direct visualization of very large images.

*Particle rendering*, where we directly visualize the particles of the APR as glyphs (see Figure 3 and Figure 5; Figure 6D and SVideo 4&5 show additional examples of particle renderings in 3D using the open-source 3D visualization software *Scenery* (*37*)).

For more details on visualization, see SuppMat 22.5.

### Image Processing Summary

Across all benchmarks and exemplar datasets, other than the worst-case example of linear neighbor access, processing directly on the APR resulted in lower execution times and memory costs. In most cases, the reductions are directly proportional to the computational ratio (CR), hence fulfilling RC4. Moreover, in the examples of visualization and segmentation, the memory cost reduction of the APR enabled processing of data sets that would not otherwise have been possible on our benchmark machine.

The APR has a range of interpretations that align with those of pixel images, allowing direct application of established image-analysis frameworks to the APR. For algorithms that require a locally isotropic neighborhood, the anisotropic local neighborhood of the APR graph can be avoided by using a particle-wise isotropic patch reconstruction, enabling also these algorithms to directly run on the APR. In many cases, however, the additional information about the structure of the image provided by the APR can be leveraged in existing algorithms, as shown here for segmentation, and it can be used to design novel algorithms, such as content-adaptive filters, variational models, and Laplacian graph-based image processing methods.

When designing novel APR-based algorithms, it may be important to account for the noise distribution (*39*). As expected, the noise distribution on the particles is different from that of the original pixels, transformed by the interpolation scheme used to compute particle intensities, and it naturally decomposes by resolution level (see SuppMat 7.6). Therefore, noise terms or regularizers in image-processing models may have to be adjusted or designed accordingly. However, the noise distribution in content-rich areas, notably around edges in the images, is largely unchanged. For image-analysis methods that focus on these areas, such as segmentation methods, the same noise models as on pixels may thus still be used.

## Discussion and Conclusion

We have introduced a novel content-adaptive image representation for fluorescence microscopy, the Adaptive Particle Representation (APR). The APR is inspired by how the human visual system effectively avoids the data and processing bottlenecks that plague modern fluorescence microscopy, particularly for 3D imaging. The APR combines aspects of previous adaptive-resolution methods, including wavelets, super-pixels, and equidistribution principles in a way that fulfills all Representation Criteria (RC) set out in the introduction. The APR is computationally efficient, suited for real-time applications at acquisition speed, and easy to implement.

We presented the ideas and concepts of the APR. The APR resamples an image by adapting a set of particles to the content of an image, taking into account the Local Intensity Scale, similar to gain control in the human visual system. The main theoretical and algorithmic contribution that made this possible with a computational cost that scales linearly with image contents is the Pulling Scheme. The Pulling Scheme guarantees image representations within user-specified relative intensity deviations.

We verified accuracy and performance of the APR using synthetic benchmark images. The analysis showed that all theoretical results hold in practice, and that the number of particles used by the APR scales with image content while maintaining image quality (RC1). Further, we showed that although image noise places a limit on representation accuracy, there exists an optimal range for the relative error threshold. In this range, the reconstruction error for noisy images is always well within the imaging noise level (RC1). Moreover, we found that the number of particles is independent of the original image size, with computational and memory costs of the APR instead proportional to the information content of the image (RC2). We showed how pixel images can rapidly be transformed to the APR, and efficiently stored both in memory and in files (RC3). We have demonstrated that the APR benefits, both in terms of execution time and memory requirements, can be leveraged for a range of image-processing tasks without ever returning to a pixel image, with minimal changes to existing algorithms (RC4). Finally, we have shown how the APR enables the development of novel, content-adaptive image-processing algorithms.

Taken together, the APR meets all four representation criteria set out in the introduction. We believe that the gains of the APR will in many cases be sufficient to alleviate the current processing bottlenecks. In particular, image-processing pipelines using the APR would be well suited for high-throughput experiments and real-time processing, e.g., in smart microscopes (*9,40*). However, the APR is sub-optimal with respect to the number of particles used. This sub-optimality results from the conservative assumptions required to derive the efficient Pulling Scheme, and from the generality of the Reconstruction Condition. It is proven by the fact that the APR particle properties can be represented by a Haar wavelet transform (*17*) with a number of non-zero coefficients that is either equal to or less than the number of particles in the APR, while allowing exact reconstruction of the APR particle properties (SuppMat 12).

The use of adaptive representations of images (*22–24*) and its motivation by the human visual system (*13, 18*) are not new. While the adaptive placement of the particles in the APR bears visual similarity to half-toning methods and techniques based on the Floyd-Steinberg error-diffusion algorithm (*41*), the mathematical foundations and the algorithms themselves differ fundamentally. The APR does, however, share several concepts with established adaptive representations. The Resolution Function *R*(*y*) of the APR, e.g., is related to the oracle adaptive regression method (*42*), and the derivation and form of the Resolution Bound are related to ideas originally introduced in equi-distribution methods for splines (*43–45*), which also inspired the work here (*46*). The Reconstruction Condition for a constant Local Intensity Scale relates to infinity-norm adaptation (*47*) for wavelet thresholding as used in adaptive surface representations. Finally, powers-of-two decomposition of the domain is central to many adaptive-resolution methods (*17, 19, 36, 48*) and its use here was particularly inspired by Ref. (*49*). Despite these relations to existing methods, the APR uniquely fulfills all representation criteria and extends or links many of the previous concepts.

Core novelties of the APR include the spatially varying Local Intensity Scale, the broad class of reconstruction methods available, “backwards compatibility” to pixel images by on-the-fly local patch reconstruction, guaranteed theoretical bounds on the representation accuracy, and the ability to combine existing compression schemes with the APR. Moreover, the computational efficiency of the APR enables real-time workflows where images are transformed at acquisition rate.

## Outlook

The APR has the potential to replace pixel-based image-processing pipelines for the next generation of fluorescence microscopes. We envision that the APR is immediately formed, possibly after image enhancement (*50*), on the acquisition computer or even on the camera itself. Following this, all data transfer, storage, visualization, and processing can be done on the APR, relaxing downstream bottlenecks. In cases where regulatory requirements or statistical noise analyses require the raw pixel data to be archived, this is best done by archiving the difference image between the raw pixel data and the APR. Since the APR captures all imaged structures, the difference image is typically very sparse and can effectively be compressed. All processing can then still be done on the APR from which the raw pixel data can exactly be reconstructed using the archived difference image whenever needed.

The realization of APR-based pipelines requires further algorithm and software development, including GPU acceleration and integration with current microscope systems, image databases (*51*), and image-processing tools (*52*). This is achieved through wrappers of the provided C++ Library *LibAPR* (github.com/cheesema/LibAPR).

Here, we presented a particular realization of an APR pipeline. We foresee alternative pipelines, e.g., using deep learning approaches (*53*) to provide improved estimation of the Local Intensity Scale, the image intensity gradient, and the smooth image reconstruction. Just as in space, the APR can also be used to adaptively sample time. Such temporal adaptation can lead to a multiplicative reduction in memory and computational costs compared to those presented here. Further, the APR can be extended to allow for anisotropic adaptation using rectangular particle cells, local affine transformations, or anisotropic particle distributions within each cell.

Given the wide success of adaptive representations in scientific computing, the unique features of the APR could be useful also in non-imaging applications. This includes applications to time-series data, where the APR could provide an adaptive regression method (*42*), to surface representation in computer graphics (*47*), and to numerically solving partial differential equations with spatial adaptivity (*46, 54–56*).

## Acknowledgements

We would like to thank the members and leaders of the Tomancak Lab at the Max Planck Institute of Molecular Cell Biology and Genetics (MPI-CBG), Huisken Lab at the MPI-CBG and Morgridge Institute for Research, Royer Lab at the CZ Biohub, Keller Lab at Janelia Farm, Lemaire Lab at Centre de Recherches de Biochimie Macromolculaire, and the Deutsches Zentrum fr Neurodegenerative Erkrankungen e.V. for the generous allowance of the use of their datasets during the bench-marking and development of this work. Further, we would like to thank Michael Hecht for discussions regarding notation, and Jan Huisken for his feedback during the development of the APR.

## Footnotes

1 Alternative implementations are possible that do not require the explicit storage of the full tree structure, but are not discussed here.

2 MCR = (Size of the input image file in Bytes)/(Size of the compressed APR file in Bytes)

